# Deciphering the rule of antigen-antibody amino acid interaction

**DOI:** 10.1101/2023.05.05.539546

**Authors:** Min Jiang, Changyin Fang, Yongping Ma

## Abstract

Antigenic drift is the biggest challenge for mutagenic RNA virus vaccine development. The most fundamental but neglected thing is to determine the immune escape mutation map (IEMM) of 20 amino acids to reveal the rule of the viral immune escape. We use universal protein tags as a linear epitope model to determine the relationship between the epitope mutation and immune escape. To describe and draw amino acid interaction maps, mutations of protein tags are classified into four types: IEM (immune escape mutation), ADERM (antibody-dependent enhancement risk mutation), EQM (equivalent mutation), and IVM (invalid mutation). To make up for the data limitation, the amino acid interaction pairs determined by 3D structure through literature search are simultaneously used to form a more systematic and universal antigen-antibody interaction maps. The results are: (i) one residue interacts with multiple amino acids in antigen-antibody interaction; (ii) Most amino acid replacements are IVM and EQM. (iii) Once aromatic amino acids replace non-aromatic amino acids, the mutation is often inactive. (iv) Substituting residues with the same physical and chemical properties easily lead to IEM. Therefore, this study had important theoretical significance for future research on antigenic drift, antibody rescue and vaccine renewal design.

**Importance:** We typed the antigenic epitope mutations into IEM, ADERM, EQM, and IVM types to describe and quantify the results of antigenic mutations. According to the results, the antigen-antibody interaction rule was summarized as one-to-many interaction rule. To sum up, the Epitope mutation rules were defined as IVM and EQM predomination rule, aryl mutation escape rule and homogeneous mutation escape rule.

## INTRODUCTION

Because RNA polymerase lacks the error-correcting mechanism of 5’-3’ exonuclease and causes the genetic variation of the virus *(1)*. When this mutation produces amino acid substitution in the neutralizing antigen (Ag), which lead to typical antigen drift and immune escape (2). Once the immunodominant epitope of the virus surface protein is mutated to form a new subtype, the existing neutralizing antibody (Ab) no longer neutralizes the mutated virus (3). This phenomenon occurs repeatedly in the severe acute respiratory syndrome coronavirus 2 (SARS-CoV-2) epidemic (4-6). SARS-CoV-2 is the pathogen of novel coronavirus pneumonia named COVID-19, resulting in significant morbidity and mortality (7, 8).

Like most mutagenic RNA viruses, SARS-CoV-2 has high genetic variability and rapid evolution *(9, 10)*. Because the SARS-CoV-2 mutants in the current epidemic are resistant to neutralizing Abs, how to solve antigenic drift is a substantial theoretical and practical problem (4). The strains B.1.617.2 and B.1.1.529 have swept the world and led the virus to evade the immune response (11-15). This has forced the redesign and production of new vaccines to cope with the new variants (16). However, the dilemma is that vaccine development cannot keep pace with viral mutations.

Antigenic drift is the typical process of RNA virus. According to the antigenic drift, successfully model, new strains may be generated constantly through mutation. Still, most of these cannot expand in the host population due to pre-existing immune responses against epitopes of limited variability (17). Notably, this natural selection is biased, *e.g*., the mutation frequency of E484K is 5.5-fold that of E484Q (18).

Immune recognition occurs in matching and anastomosis between specific positions and specific fragments of Ab and Ag molecules. Due to the complex spatial structure of proteins and the diversity of organisms, it is very difficult to predict exactly how the antigen-determined amino acid will mutate. Anyhow, exploring the rule of amino acid interaction between Ags and Abs is of great significance, and then summarizing the interaction (recognition and binding) rule of amino acids for current virus immunity and vaccine preparation. The best way is to detect changes in the ability of the antigen to bind to the mAb by mutating the key amino acid on the epitope to summarize the regular amino acid interaction spectrum.

To describe the relationship between linear epitope mutation and immune escape, we cautiously assumed four concepts: (i) Immune escape mutation (IEM) meant that the residue substitution caused the antigen to lose its affinity (recognition) to pre-existing Ab, or to remain less than the assumed 30% affinity without neutralization. (ii) Antibody-dependent enhancement risk mutation (ADERM) referred to the residue substitution caused the antigen to remain pre-existing Ab affinity more than the assumed 30% but less than 50%. However, the pre-existing Abs could not neutralize the mutated antigen (pathogen). Rather, the virus-Ab complex with low affinity enhanced virus uptake resulting from the attachment of immune complexes to the Fcγ receptor and enhanced the infection. (iii) Equivalent mutation (EQM) meant that the residue substitution led the Ag to remain pre-existing Ab affinity beyond the assumed 50% but less than 80%. Fortunately, the pre-existing Ab still completely neutralizes the antigen. (iv) Invalid mutation (IVM) indicated that the residue substitution did not affect the pre-existing Ab affinity and the Ab completely neutralized the Ag.

The enzyme-linked immunosorbent assay (ELISA) was employed to measure the degree of binding affinity between anti-protein tags mAbs and protein tag mutants. To make up for the data limitation, we also summarized the Ag-mAb interaction pairs determined by 3D structure through a literature search.

## RESULTS

### Results of protein-tags ELISA

We screened the key residues in HA-tag to mAb TA180128. At first, HA-tag (Y1P2Y3D4V5P6D7Y8A9) was substituted one by one from Y1 to A9 with G, E, or H, respectively. Except for A9 residue, mAb TA180128 reacted to the other residues from Y1 to Y8. After substituting with other 19 amino acids from Y1 to Y8, the 15 mutants were classified as IVM (Y1H, Y1E, Y3H, V5A, V5I, V5K, V5L, V5M, V5S, V5T, D7G, D7H, Y8E, Y8G, and Y8H) for maintaining more than 80% affinity to mAb TA180128. 33 mutants were IEM (P2E, P2H, D4E, D4H, V5F, V5G, Y1G, Y3E, Y3G, and P6G) for less than 18% affinity, or IEM (P2G, V5D, V5E, V5W, V5Y, P6A, P6C, P6D, P6E, P6F, P6H, P6I, P6K, P6L, P6M, P6N, P6Q, P6R, P6S, P6T, P6V, P6W, and P6Y) for losing the affinity to mAb TA180128 completely. However, V5C, V5N, and V5R were ADERM for maintaining about 35%-47.8% affinity. Moreover, D4G, V5H, V5P, V5P, V5Q, and D7E were EQM for maintaining about 66%-79% affinity (Figure 1A).

**Figure 1.**
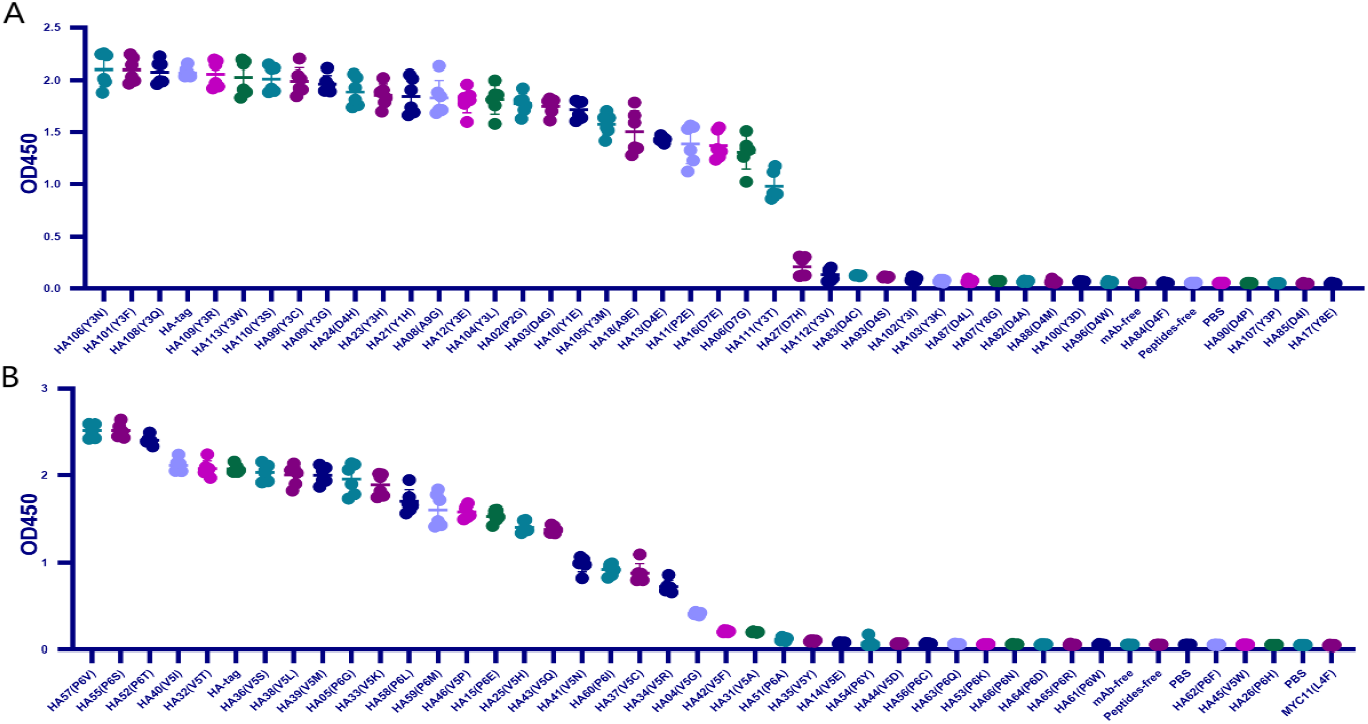
Scanning the binding affinity of HA-tag mutants to mAb TA180128. (A.B.) HA-tag mutants P2X, Y3X, D4X, V5X, P6X, D7X, Y8X, A9X.

Since IVM and EQM did not affect the protective effect of the current vaccine, the aim was to reveal the IEM of V5 and P6 substitutions replaced by the other 19 amino acids, respectively. In summary, V5 IEM substitutions accounted for 36.8% of the 19 amino acids. EQM substitutions accounted for 15.8%. More than 36.8% V5 substitutions were completely IVM and about 15% were ADERM. Similarly, P6 IEM accounted for 57.8%; EQM accounted for 10.5%; about 26.3% were IVM and 5% were ADERM (Figure 1B). Therefore, about half of antigenic drift (IVM and EQM) did not affect the immunity of mAb TA180128.

To cover 20 amino acids substitution, we added the MYC-tag, Flag-tag, and VSV-tag to the experimental catalogue. Specifically, we replaced the Q2, S6, and E8 residues of MYC-tag with G and its L4 and I5 with other 19 amino acids, respectively. While other 19 amino acids replaced the K3 of Flag-tag. G replaced the N7, R8, and I4 in VSV-tag and the T2 and M6 were mutated to the other 19 amino acids, respectively.

For c-MYC-tag mutants, except for the affinity of Q2G, I5S, and E8G > 80%, which belonged to IVM. I5L, S6G, and L4C belonged to EQM. The other mutants had low affinity or could not be combined and classified as IEM (Figure 2A).

**Figure 2.**
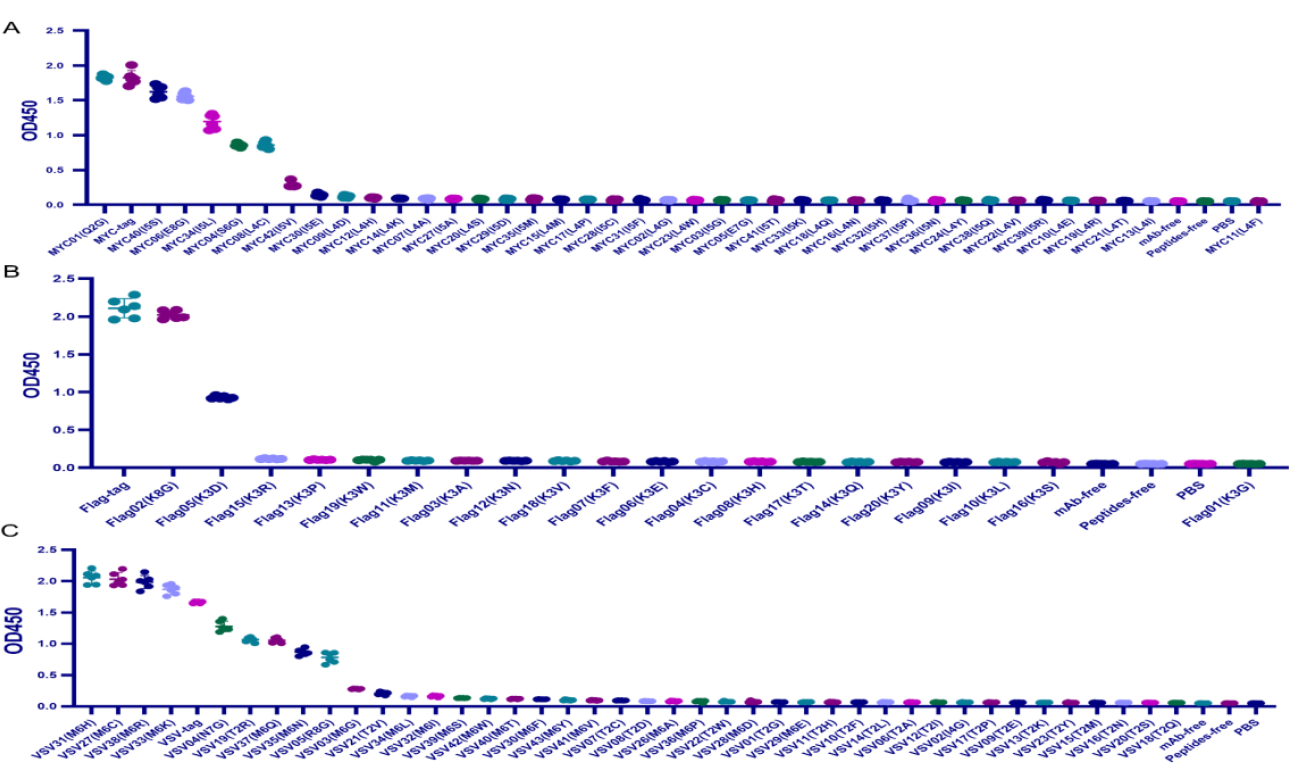
Scanning the binding affinity of MYC-tag, Flag-tag and VSV-tag mutants. (A) Scanning the binding affinity of MYC-tag mutants of Q2X, L4X, I5X, S6X, and E8X to mAb6003-2-lg. (B) Scanning the binding affinity of Flag-tag mutants K3X to mAb 66008-3-lg. Most of the replacement were IEM except for K3G and K3D. (C) Scanning the binding affinity of VSV-tag mutants of T2X, M6X, N7X, R8X, and I4X to mAb A02180.

Among the mutants of Flag-tag, only K8G shared similar mAb binding affinity with Flag-tag and was defined as EQM. The relationship of K3D was 43% and belonged to ADERM. Anyway, the other seventeen K3 mutants were IEM (Figure 2B).

Similarly, most VSV-tag mutants had low affinity as IEM. Unexpectedly, the level of affinity of the four mutants replaced by M6C, M6H, M6K, and M6R was higher than that of the VSV-tag, and the affinity of N7G, T2R and M6Q was EQM for 66%-79% affinity. However, M6N and R8G were ADERM for less than 50% affinity (Figure 2C).

To display the mutant interaction visually, we used Microsoft Excel to draw the heatmap of amino acid residue substitution. From the selected and substituted nine amino acids, I, K, L, and T tend to lose their affinity with mAb after substituting them with the other 19 amino acids (Figure 3). Except that the binding affinity substituted by C, H, K, and R could be maintained, substituting M by other amino acids led to IEM. Similarly, when E, H, and G replaced D, it could still recognize and bind to mAb. The other substitutions of D led to IEM. Conversely, about 84% of substitutions of V and Y showed IVM. Most substitutions of V and Y retained their original immune affinity. Only V/F, V/G, V/W, V/Y, Y/V, Y/D, Y/I, Y/P, and Y/K replacement models produced IEM (Figure 3). Nevertheless, only 42% of substitutions of P showed IVM, and the other 11 substitutions were IEM (Figure 3). Notably, all amino acids replaced by A, D, F, P, W, and Y led to IEM, and 98% of amino acids replaced by V and I might be IEM (Figure 3). However, 66% of amino acids replaced by G might be IEM (Figure 4).

**Figure 3.**
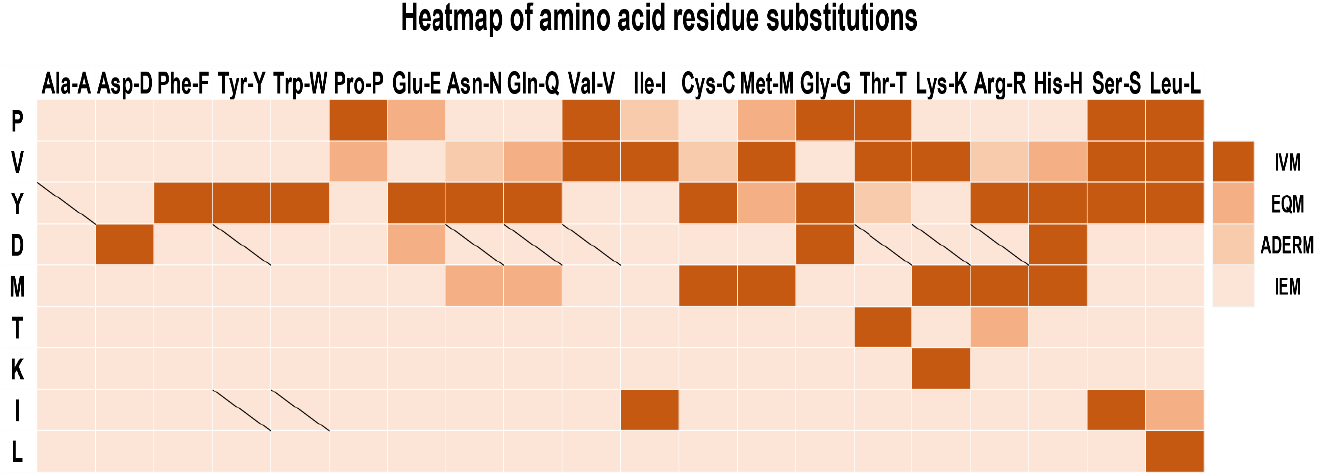
Heat map of amino acid residue substitution verified by ELISA. IVM: Invalid mutation, EQM: Equivalent mutation, ADERM: antibody-dependent enhancement risk mutation; IEM: Immune escape mutation; P: HA-P6; V: HA-V5; Y: HA-Y3; D: HA-D4; M: VSV-M6; T: VSV-T2; K: Flag-K3; I: MYC-I5; L: MYC-L4;Slash line: No experimental data available.

**Figure 4.**
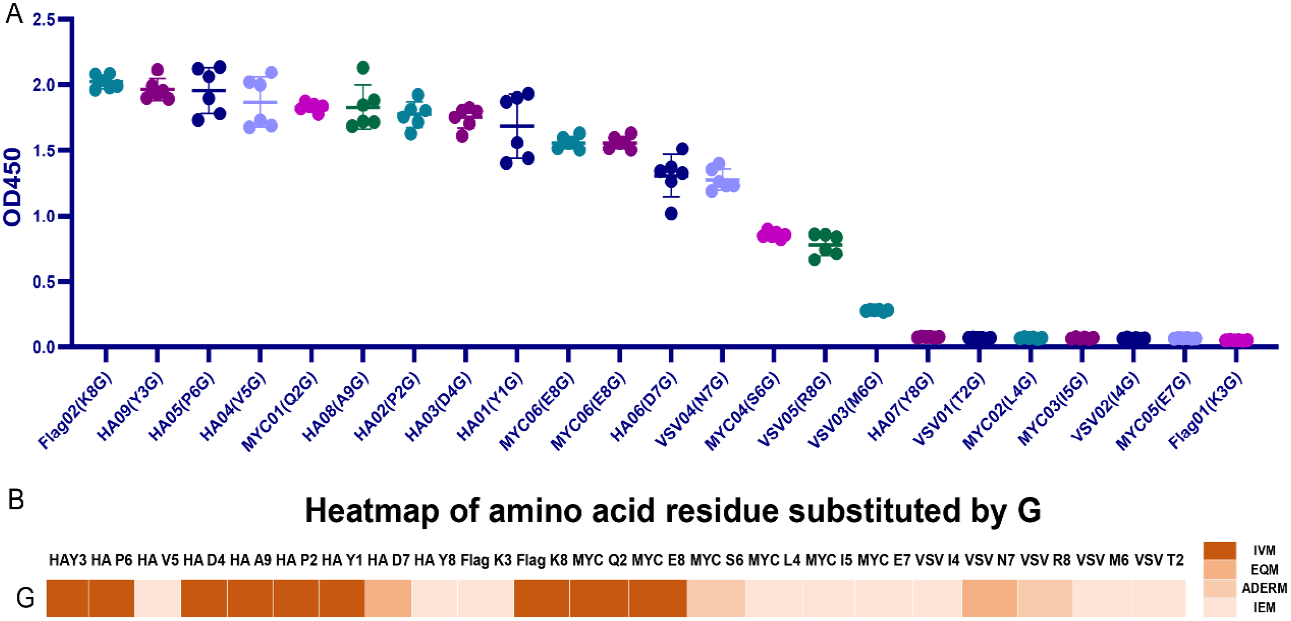
The binding affinity map of mutant tags replaced with G. (A) Map of ELISA results of mutant strains and monoclonal antibodies. (B) Heatmap of binding affinity of G replaced mutant tags and monoclonal antibodies.

### Literature data analysis of antigen-antibody interactions

Seventy articles related to Ag-Ab interaction were screened through the NCBI PubMed and Web of Science. The interactions of amino acid residues at the Ag-Ab binding interface were extracted from the 3D structures reported in the literature, and the amino acid binding matrix was drawn and colored according to the number of relevant literature (Figure 5A). The X-axis is the amino acids of the Ab, and the Y-axis is antigenic residues. According to the matrix, the binding rules of amino acids in Ags and Abs tended to be consistent. Among the 20 types of amino acids, Y, S, D, and R had an affinity with other amino acids and could recognize almost all amino acids in the literature. Instead, C and M were practically silent and only interacted with relatively few amino acids, coinciding with some of our ELISA experiments’ findings. Explicitly, residue M did not find in the Ab interface and only 4 in the Ag interface (Figures 5B and C). Residue C found only one in the Ab interface and only 3 in the Ag interface (Figures 5B and C). Otherwise, P was rarely found in Ab interfaces and V was rarely in Ag interfaces (Figures 5B and C).

**Figure 5.**
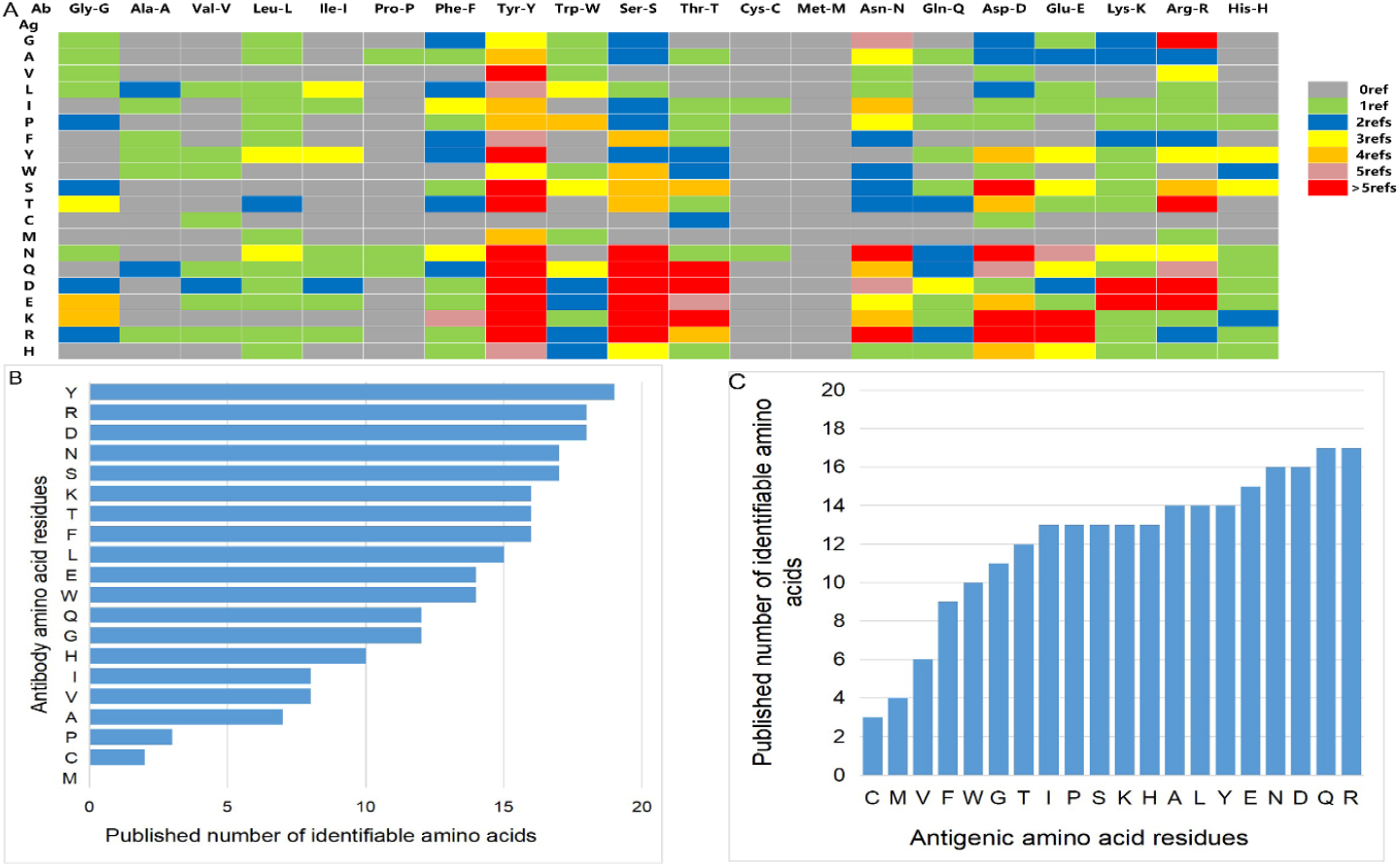
Summary of literature amino acid recognition rules. (A) Amino acid recognition rule matrix collected by literature. (B) Histogram of identifiable amount of amino acid residues in antibodies. (C) Histogram of identifiable quantity of amino acid residues in antigen.

Next, the histogram of the interaction capacity of the 20 amino acids was plotted according to the literature. The top 10 amino acids with the highest Ab detection rate were Y, R, D, N, S, K, T, F, L, and E, respectively (Figure 4B). However, the top 10 detection amino acids in Ags were R, Q, D, N, E, Y, L, A, H, K, S, P, S, and I (Figure 5C). It must explain that the H, K, S, P, S, and I shared the 10th rank for their same detection rate (Figure 5C).

### Amino acid interaction network analysis

The data were imported into Cytoscape to draw an interaction network. According to the physical and chemical properties, 20 amino acids were divided into 8 categories, which were aliphatic amino acids, aromatic amino acids, acidic amino acids, alkaline amino acids, amide amino acids, sulfur-containing amino acids, hydroxyl amino acids, and imino acids. Then, the interaction diagram was constructed (Figure 6A). Except for a few aliphatic amino acids and all sulfur-containing amino acids, the side chains of hydroxy-amino acids S and T could interact with 14 amino acids in the literature (Figure 6B). The aliphatic amino acids included A, V, I, and L, and only I could interact with C (Figure 6C). The alkaline amino acids K, R, and H could interact with 17 kinds of amino acids in Abs (Figure 6D). Notably, antigenic P interacted with 11 amino acids in Abs, and P in Abs only interacted with antigenic N, Q, and R (Figures 6A, C, E, and F). The antigenic C (interacted with D, T, and V) and M (interacted with L, R, W, and Y) contained sulphur in their side chains, causing them not to cross-link (Figure 6G). Otherwise, the residue of Ab C only recognized antigenic I and N (Figures 6A, C, and F). Acidic amino acids could interact with 16 amino acids (Figure 6H). Similarly, antigenic aromatic amino acids seemed to interact with more than 10 amino acids (Figure 6I). To quickly view the amino acid interaction between Ag-Ab, single interaction network was plotted individually in alphabetical order of amino acid abbreviations. Amino acids without interaction were listed below the main figure in boldface (Figure 7).

**Figure 6.**
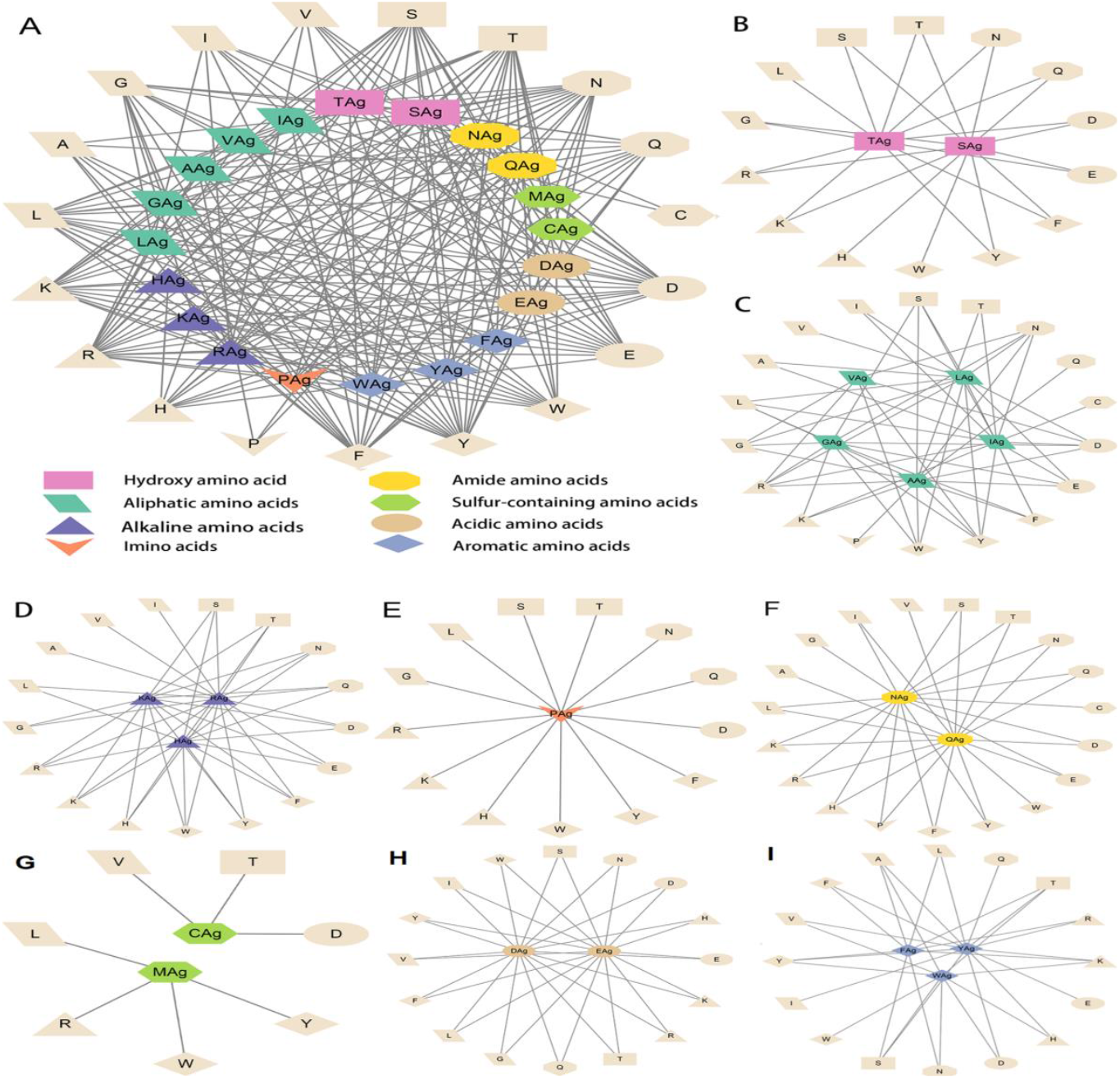
Literature antigen-antibody amino acid interaction diagram. (A) Summary diagram of Ag-Ab amino acid interaction based on physical and chemical properties. (B) Diagram of interactions between hydroxy amino acids and 20 amino acids. (C) Diagram of interactions between aliphatic amino acids and 20 amino acids. (D) Diagram of interactions between alkaline amino acids and 20 amino acids. (E) Imino acids. (F) Amide amino acids. (G) Sulfur-containing amino acids. (H) Acidic amino acids. (I) Aromatic amino acids. The antigenic amino acid abbreviation was labeled with the subscript Ag to distinguish antibody amino acids.

**Figure 7.**
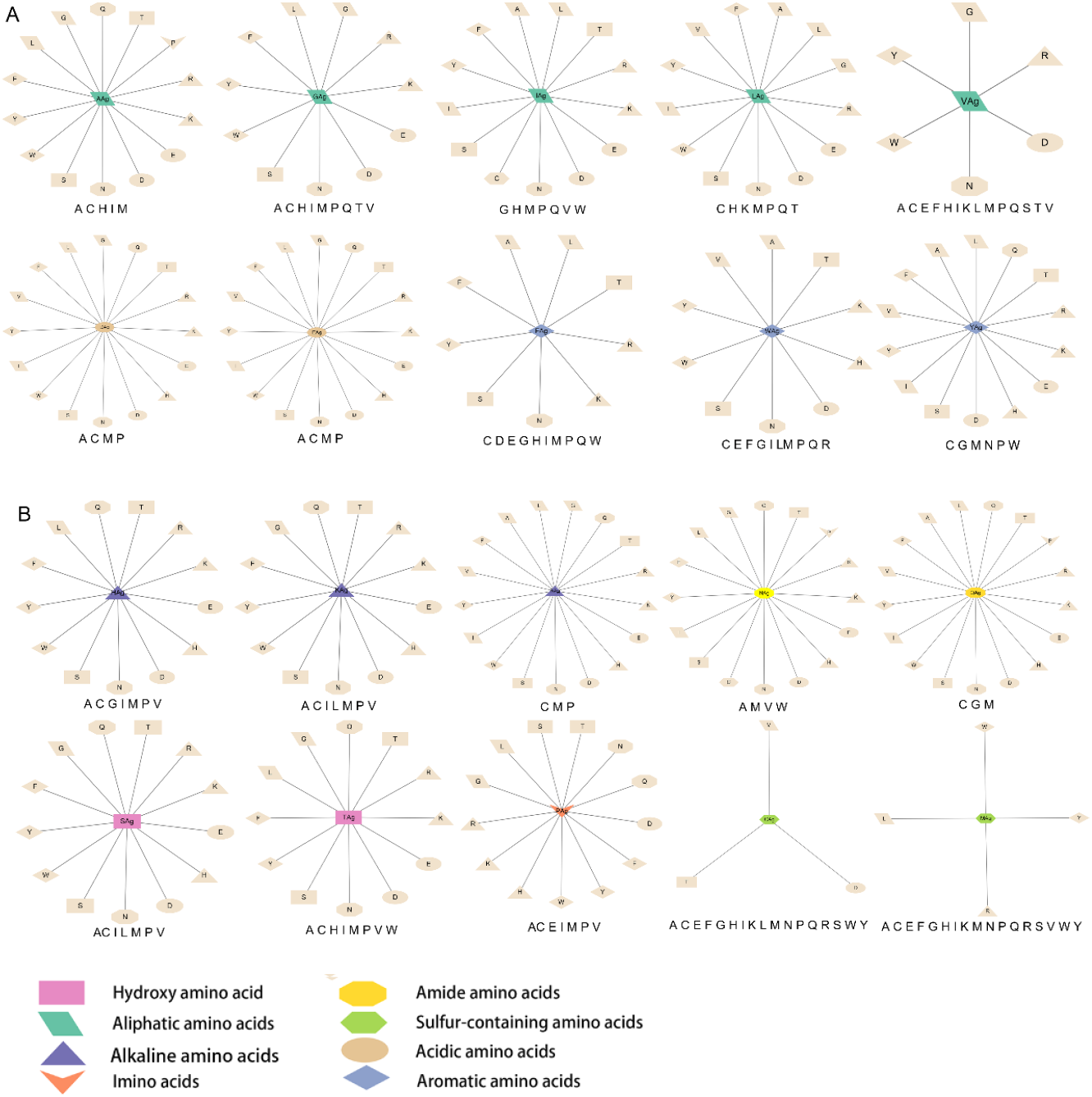
20 antigenic amino acids interaction plotted individually. Antibody amino acids without interaction were listed in boxes below the main figure in bold type. The antigenic amino acid abbreviation was labeled with the subscript Ag to distinguish antibody amino acids.

### Verification of experimental data and literature data

To reveal the effect of mutation on the peptide structure and verify the experimental results, we used the computer ab initio folding algorithm to predict the 3D structure of peptide tags and their mutants. We predicted the 3D structure of HA-tag, MYC-tag, and Flag-tag and their mutants (HA-Y1G, HA-Y3G, HA-Y8G, MYC-E8G, MYC-E7G, Flag-K8G, and Flag-K3G), respectively, by online tool PEP-FOLD3. After obtaining the PDB file, we compared the mutant with the original structure. The fit degree of HA-Y8G and MYC-E7G to the original tag structure was lower than other mutants (Figure 8). Surprisingly, the Flag-K3G mutation disrupted the original α-helix and curled into a semi-O-ring in the opposite direction of the original Flag-K3 peptide, affecting Ab binding capacity (Figures 2B and S2). Consequently, the low fit mutation resulted in IEM due to loss of interaction. The same IEM mutations were happened in HA-D4E, HA-P6F, HA-P6W, HA-P6Y, VSV-M6F, VSV-M6Y, and VSV-M6W (Figures 9 and 10).

**Figure 8.**
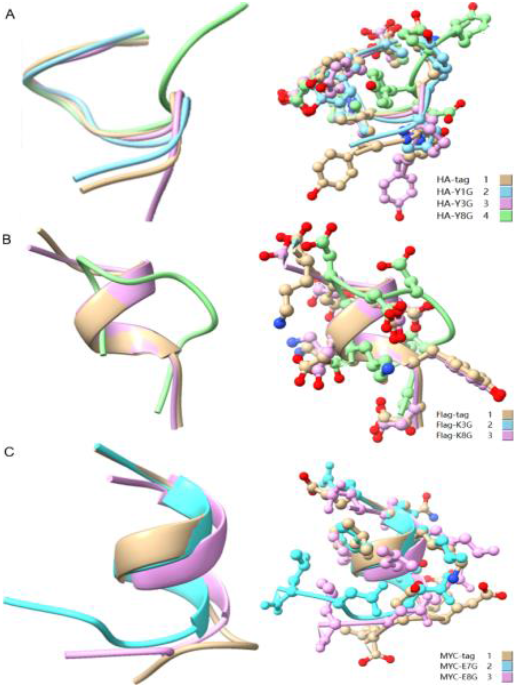
Comparison of the 3D structure of mutant tags replaced with G. (A) HA-tag with HA-Y1G, HA-Y3G and HA-Y8G (B) Flag-tag with Flag-K8G and Flag-K3G (C) MYC-tag with MYC-E8G and MYC-E7G.

**Figure 9.**
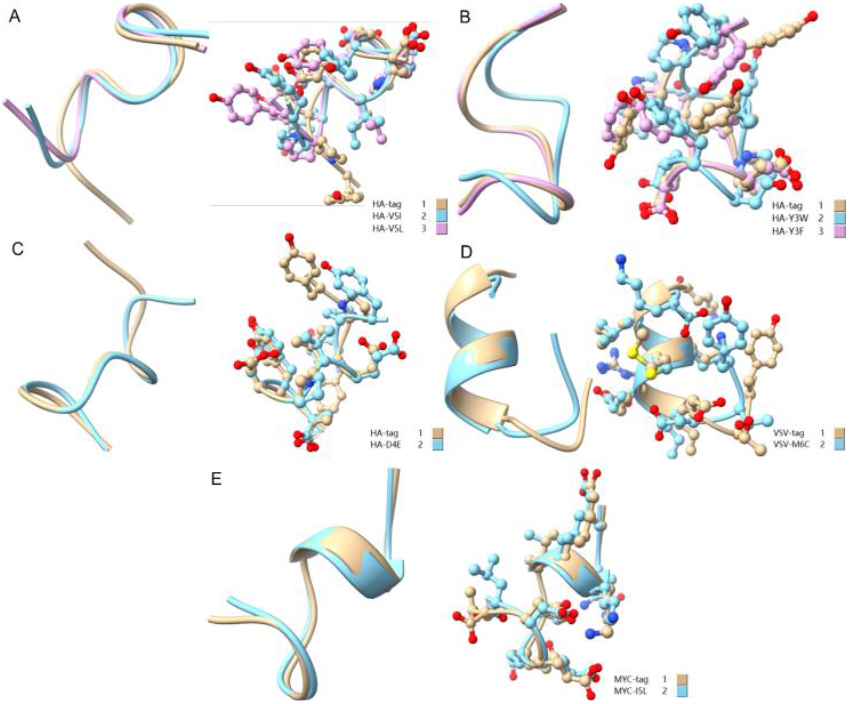
Comparison of amino acid mutant with similar physicochemical property residues. (A) HA - tag, HA - V5I, and HA - V5L; (B) HA-tag, HA-Y3W, and HA-Y3F; (C) HA-tag and HA-D4E; (D) VSV-tag and VSV-M6C; (E) MYC-tag and MYC-I5L.

**Figure 10.**
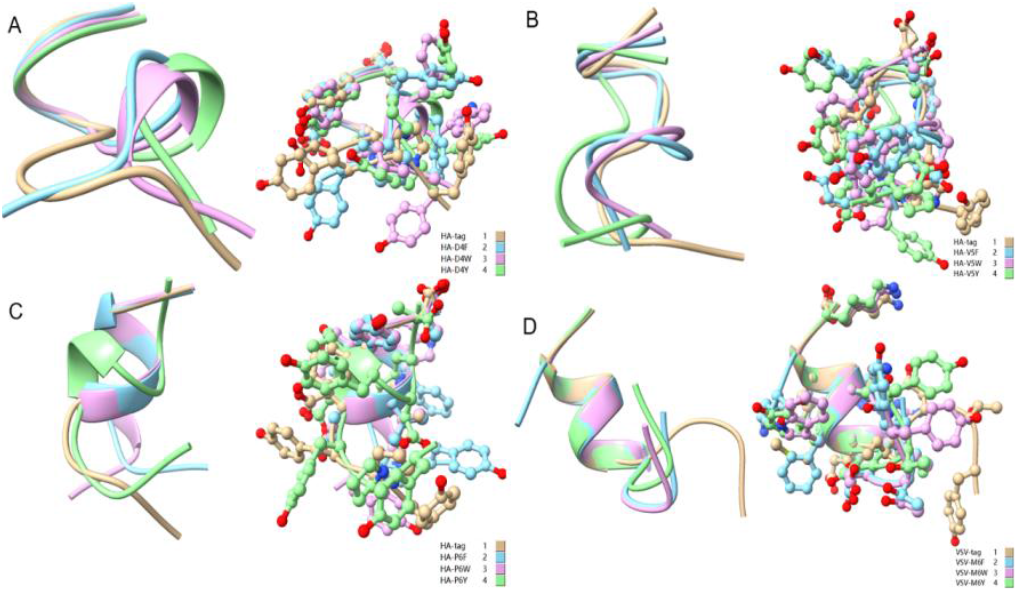
Comparison of aromatic amino acid substitution mutants. (A) HA-tag, HA-D4F, HA-D4Y, and HA-D4W; (B) HA-tag, HA-V5F, HA-V5Y, and HA-V5W; (C) HA-tag, HA-P6F, HA-P6Y, and HA-P6W((D) VSV-tag, VSV-M6F, VSV-M6Y, and VSV-M6W.

Protein tags mutation experiment found that the substitution of amino acids with similar physical and chemical properties residue tended to IVM, *e.g*., aliphatic amino acids V, I, and L substituted each other (HA-V5I, HA-V5L; MYC-I5L) (Figures 1, 2A and 2C). Similarly, aromatic amino acids Y, W, and F mutual substitution were also IVM, *e.g*., HA-Y3F and HA-Y3W (Figure 1). The results indicated that the amino acids with the same physical and chemical properties of the side chain group shared high similarity in 3D structures (Figure 9). In addition, the non-aromatic amino acids were replaced by aromatic amino acids (F, Y, and W) and often lost their affinity to antibodies (Figures 1 and 2). It might be related to the benzene ring on the side chain of aromatic amino acids. We consequently predicted the structure of some non-aromatic amino acid mutants replaced by aromatic amino acids and compared their structural changes (Figure 10). To visually describe the structural changes of the mutated tags, the RMSD values were obtained in UCSF ChimeraX for the fitted structures of the above-mentioned mutants and the original protein tags were summarized (Table 1). If RMSD was less than 3 Å, the two structures were considered similar. Conversely, if the RMSD was more significant than 3 Å, the two structures did not match. Thus, our test suggested that the protein-tag mutants with RMSD > 3.0 Å were trended to IEM (Table 1).

**Table 1.** RMSD of the fitted structures of mutant tags and original protein-tags.

| Mutant | RMSD(Å) |
| --- | --- |
| HA-Y1G | 0.776 |
| HA-Y3G | 1.594 |
| HA-Y8G | 4.496 |
| MYC-E8G | 2.656 |
| MYC-E7G | 3.835 |
| Flag-K3G | 4.682 |
| Flag-K8G | 0.408 |
| HA-V5L | 1.825 |
| HA-V5I | 1.885 |
| HA-Y3F | 0.332 |
| HA-Y3W | 2.457 |
| HA-D4E | 1.845 |
| VSV-M6C | 2.799 |
| MYC-I5L | 0.557 |
| HA-D4F | 4.139 |
| HA-D4W | 3.401 |
| HA-D4Y | 4.301 |
| HA-P6F | 3.792 |
| HA-P6W | 4.189 |
| HA-P6Y | 4.214 |
| HA-V5F | 4.760 |
| HA-V5W | 3.509 |
| HA-V5Y | 3.650 |
| VSV-M6F | 4.550 |
| VSV-M6W | 4.212 |
| VSV-M6Y | 4.175 |

RMSD < 3.0 referred higher goodness of fit.

Therefore, this study had important theoretical significance for future research of antigenic drift and rescue mAb. Moreover, the following conclusions were summarized:

I. Ag-Ab interactions were classified into four types: IEM, ADERM, EQM, and IVM, according to antibody binding affinity.
II. One-to-many interaction rule: For most antibody and antigenic amino acids, the pattern of recognition and binding was one-to-many mode. The conclusion was verified by the literature data.
III. IVM and EQM predomination rule: Most of the replacements for specific antigenic amino acids were IVM and EQM.
IV. Aryl mutation escape rule: Once aromatic amino acids replaced an antigenic non-aromatic amino acid, the mutation was often inactive and resulted in IEM.
V. Homogeneous mutation escape rule: Most replacement mutations easily led to IEM when other amino acids replaced antigenic amino acids with similar physical and chemical properties.

## DISCUSSION

Viral immune escape caused by antigenic drift has always been a great challenge for vaccination and prevention (19). This study aims to establish the rule of the amino acid interaction during Ag-Ab recognition and use this rule to guide scientists to update vaccine or rescue ineffective mAb due to antigenic mutations in clinical applications. Ab rescue refers to when an essential antigenic amino acid mutation causes the failure of mAb in clinical application, the original amino acid on the mAb is replaced with a new amino acid according to the Ag-Ab recognition rule to recognize and neutralize the mutated antigen again. We termed it the reverse antibody technique. For instance, the E484K mutation of SARS-CoV-2 RBD disabled mAb P17 neutralization (20, 21). 3D data demonstrated that the negatively charged E484 interacted with positively charged H35, H99, and R96 residues, which formed a strong electrostatic interaction (21). Thus, the E484K mutation was repulsive to the positively charged binding site of the mAb P17. Guided by our reverse antibody theory, mAb P17 might be theoretically rescued by mutating part or all of the H35, H99, and R96 into negatively charged residues such as E and/or D.

The IEM rule obtained from protein tags was confirmed in the real-world data. For example, HA-tag P6L mutation resulted in IEM (Figure 1). In SARS-CoV, the P462L mutation led to escape from mAb CR3014 (22). In addition, the spike L452R mutation conferred SARS-CoV-2 escape from the immune (23). Our findings suggested that Ags were often not recognized by Ab when other residues replaced antigenic residues K, L, or T. Correspondingly, the A180V mutation in the influenza virus promotes the virus to escape from Ab-based immunity (24-26). V483A mutation in SARS-CoV-2 was resistant to some neutralizing antibodies (27). In this study, the mutations of V to A led to IEM (Figure 1). The escape caused by these mutations is consistent with the escape rules we found in this study. The amino acid mutation type of Ag and the corresponding IEM pattern are termed immune escape mutation map (IEMM). IEMM establishment aims to deal with antigenic drift and immune escape of human viruses. However, IEMM is currently just a hypothesis we proposed and needed to prove and refine in the future.

Theoretically, substituting epitope key residues seems to be a random event, but it has its rules. Many mutations do not exist in nature for immune pressure (28). That is, all strains found to have mutations in neutralizing antigens are immune escape strains. In particular, the single nucleotide mutations of L452 in SARS-CoV-2 are limited to Q, M, and R in nature (3, 23, 29, 30). Tan et al. characterized all 19 possible mutations at L452. They revealed that five mutants (Q, K, H, M, and R) gained greater infectivity and immune escape. The other mutants failed to maintain expression or pseudovirus infectivity (31).

Although supported by experimental and literature data, our study still has some shortcomings. First, the Ag-Ab amino acids interaction pattern was not covered all 20 kinds of amino acid for deficiency of certain amino acids in linear epitopes. Even so, the joint 3D database analysis made up for this shortcoming somewhat. Second, the Ag-Ab interaction data derived from the linear protein tag epitopes may not fully reflect the true spatial conformation of complete antigen molecules. Therefore, the reverse antibody test was the main task to verify this theory in the future.

## MATERIALS AND METHODS

### Peptides and agents

Peptides of HA-tag, c-MYC-tag, VSV-tag, Flag-tag, and their mutants (more than 98% purity) were synthesized commercially by Nanjing Yuanpeptide Biotech (Nanjing, China) (Tables S2-S5). Mouse anti-HA-tag mAb were purchased from Qrigene (cat# TA180128, MD, USA). Mouse anti-c-MYC-tag and anti-Flag-tag mAb were purchased from Proteintech(cat# 6003-2-lg/66008-3-lg, Wuhan, China), Mouse anti-VSV-tag mAb were purchased from Abbkine Scientific (cat# A02180, Wuhan, China), respectively. HRP conjugated AffiniPure goat anti-mouse IgG (Proteintech, cat# SA00001-15, Wuhan, China) and DAB color development kit (cat# AR1026) were purchased from a Biological company. The other buffers were prepared in our laboratory.

### Enzyme linked immunosorbent assay (ELISA)

The peptides of tags and their mutants were diluted to 2.0 μg/ml and each was added 50 μl into 96 well plates for 1 h to adsorb (n=3). After washing with phosphate buffer saline containing 0.05% Tween-20 (PBST) three times, the plate was blocked with rapid blocking buffer for 10 min. After washing with PBST three times, mouse anti-tags mAb were added into each well (40 μl/well), and incubated at 37°C for 1 h. Next, washing and blocking as described above, 100 μl of horseradish peroxidase (HRP)-labeled goat anti-mice IgG (1:2500) were added and incubated at 37°C for 1 h. The results were detected by plate reader after color-developing with TMB single-component substrate solution (Solarbio, cat# PR1200, Beijing, China) and stopping. Peptides-free and mAb-free wells were performed as negative controls. The absorbance value at 450 nm (OD_450_) was measured for analysis.

### Literature 3D-structure references and amino acid-amino acid interaction assay

A literature search was performed using the keywords: ‘protein ‘interaction’, ‘antigen-antibody ‘interaction’, and ‘crystal structure of binding ‘surface’ in the NCBI PubMed database (https://www.ncbi.nlm.nih.gov/pubmed/) and Web of Science (https://www.webofscience.com/wos/alldb/basic-search) to collect the amino acid recognition binding data documented in the literature.

All collected data were entered into Microsoft Excel™ and visualized with Cytoscape and GraphPad Prism 8.02 software. After partial mutation, the mutant polypeptide sequence and the original protein-tag sequence of IVM and IEM were input into the online prediction software PEP-FOLD3, and the 3D structure was obtained. The best coupling model was selected according to the score. At the same time, the PDB file was downloaded to UCSF Chimera X, and the Matchmaker program was performed on the 3D structure of the mutant and the original protein tag to obtain the structure distance RMSD (root mean square deviation) value and the 3D conformational map.

## Acknowledgments

The research was funded by a research grant from the National Natural Science Foundation of China (30972585). We thanks Tayyaba Yasin for language editing.

## Additional information Declaration of competing interests

Authors declare that they have no competing interests.

## Author contributions

Y.M. conceived of the paper and written the manuscript, proposed the four concepts: IEM, IVM, EQM and ADERM, and proposed the hypothesis of reverse monoclonal antibody technology. M.J. and C.F. contributed, as co-first author, to perform the experiments, and analyze data. M.J. prepared all of the figures and written the original draft.

## Data availability

All data are available in the main text or the supplementary materials.

## Supplementary Tables

Table **S2**. Sequences of HA-tag and its mutants.

Table S3. Sequences of MYC-tag and its mutants.

Table S4. Sequences of Flag-tag and its mutants.

Table S5. Sequences of VSV-tag and its mutants.

