## Supplemental table 2-5 for "Deciphering the rule of antigen-antibody amino acid interaction"

#### Supplementary Tables

**Supplementary Table S2 Sequences of HA-tag and its mutants**

| Number | sequences |
|---|---|
| HA-tag | **YPYDVPDYA** |
| HA01(Y1G) | **GPYDVPDYA** |
| HA02(P2G) | Y**GYDVPDYA** |
| HA03(D4G) | YPY**GVPDYA** |
| HA04(V5G) | YPYD**GPDYA** |
| HA05(P6G) | YPYDV**GDYA** |
| HA06(D7G) | YPYDVP**GYA** |
| HA07(Y8G) | YPYDVPD**GA** |
| HA08(A9G) | YPYDVPDY**G** |
| HA09(Y3G) | YP**GDVPDYA** |
| HA10(Y1E) | **EPYDVPDYA** |
| HA11(P2E) | Y**EYDVPDYA** |
| HA12(Y3E) | YP**EDVPDYA** |
| HA13(D4E) | YPY**EVPDYA** |
| HA14(V5E) | YPYD**EPDYA** |
| HA15(P6E) | YPYDV**EDYA** |
| HA16(D7E) | YPYDVP**EYA** |
| HA17(Y8E) | YPYDVPD**EA** |
| HA18(A9E) | YPYDVPDY**E** |
| HA21(Y1H) | **HPYDVPDYA** |
| HA22(P2H) | Y**HYDVPDYA** |
| HA23(Y3H) | YP**HDVPDYA** |
| HA24(D4H) | YPY**HVPDYA** |
| HA25(V5H) | YPYD**HPDYA** |
| HA26(P6H) | YPYDV**HDYA** |
| HA27(D7H) | YPYDVP**HYA** |

| | |
|---|---|
| HA28(Y8H) | YPYDVPD**HA** |
| HA29(A9H) | YPYDVPDY**H** |
| HA31(V5A) | YPYD**APDYA** |
| HA32(V5T) | YPYD**TPDYA** |
| HA33(V5K) | YPYD**KPDYA** |
| HA34(V5R) | YPYD**RPDYA** |
| HA35(V5Y) | YPYD**YPDYA** |
| HA36(V5S) | YPYD**SPDYA** |
| HA37(V5C) | YPYD**CPDYA** |
| HA38(V5L) | YPYD**LPDYA** |
| HA39(V5M) | YPYD**MPDYA** |
| HA40(V5I) | YPYD**IPDYA** |
| HA41(V5N) | YPYD**NPDYA** |
| HA42(V5F) | YPYD**FPDYA** |
| HA43(V5Q) | YPYD**QPDYA** |
| HA44(V5D) | YPYD**DPDYA** |
| HA45(V5W) | YPYD**WPDYA** |
| HA46(V5P) | YPYD**PPDYA** |
| HA51(P6A) | YPYDV**ADYA** |
| HA52(P6T) | YPYDV**TDYA** |
| HA53(P6K) | YPYDV**KDYA** |
| HA54(P6Y) | YPYDV**YDYA** |
| HA55(P6S) | YPYDV**SDYA** |
| HA56(P6C) | YPYDV**CDYA** |
| HA57(P6V) | YPYDV**VDYA** |
| HA58(P6L) | YPYDV**LDYA** |
| HA59(P6M) | YPYDV**MDYA** |
| HA60(P6I) | YPYDV**IDYA** |
| HA61(P6W) | YPYDV**WDYA** |

| | |
|---|---|
| HA62(P6F) | YPYDV**FDYA** |
| HA63(P6Q) | YPYDV**QDYA** |
| HA64(P6D) | YPYDV**DDYA** |
| HA65(P6R) | YPYDV**RDYA** |
| HA66(P6N) | YPYDV**NDYA** |
| HA82(D4A) | YPY**A**VPDYA |
| HA83(D4C) | YPY**C**VPDYA |
| HA84(D4F) | YPY**F**VPDYA |
| HA85(D4I) | YPY**I**VPDYA |
| HA87(D4L) | YPY**L**VPDYA |
| HA88(D4M) | YPY**M**VPDYA |
| HA90(D4P) | YPY**P**VPDYA |
| HA93(D4S) | YPY**S**VPDYA |
| HA96(D4W) | YPY**W**VPDYA |
| HA99(Y3C) | YP**C**DVPDYA |
| HA100(Y3D) | YP**D**DVPDYA |
| HA101(Y3F) | YP**F**DVPDYA |
| HA102(Y3I) | YP**I**DVPDYA |
| HA103(Y3K) | YP**K**DVPDYA |
| HA104(Y3L) | YP**L**DVPDYA |
| HA105(Y3M) | YP**M**DVPDYA |
| HA106(Y3N) | YP**N**DVPDYA |
| HA107(Y3P) | YP**P**DVPDYA |
| HA108(Y3Q) | YP**Q**DVPDYA |
| HA109(Y3R) | YP**R**DVPDYA |
| HA110(Y3S) | YP**S**DVPDYA |
| HA111(Y3T) | YP**T**DVPDYA |
| HA112(Y3V) | YP**V**DVPDYA |
| HA113(Y3W) | YP**W**DVPDYA |

### the replaced residues were highlighted in red color.

**Supplementary Table S3 Sequences of MYC-tag and its mutants**

| **Number** | **sequences** |
|---|---|
| MYC-tag | **EQKLISEEDL** |
| MYC01(Q2G) | E**G**KLISEEDL |
| MYC02(L4G) | EQK**G**ISEEDL |
| MYC03(I5G) | EQKL**G**SEEDL |
| MYC04(S6G) | EQKLI**G**EEDL |
| MYC05(E7G) | EQKLIS**G**EDL |
| MYC06(E8G) | EQKLISE**G**DL |
| MYC07(L4A) | EQK**A**ISEEDL |
| MYC08(L4C) | EQK**C**ISEEDL |
| MYC09(L4D) | EQK**D**ISEEDL |
| MYC10(L4E) | EQK**E**ISEEDL |
| MYC12(L4H) | EQK**H**ISEEDL |
| MYC13(L4I) | EQK**I**ISEEDL |
| MYC14(L4K) | EQK**K**ISEEDL |
| MYC15(L4M) | EQK**M**ISEEDL |
| MYC16(L4N) | EQK**N**ISEEDL |
| MYC17(L4P) | EQK**P**ISEEDL |
| MYC18(L4Q) | EQK**Q**ISEEDL |
| MYC19(L4R) | EQK**R**ISEEDL |
| MYC20(L4S) | EQK**S**ISEEDL |
| MYC21(L4T) | EQK**T**ISEEDL |
| MYC22(L4V) | EQK**V**ISEEDL |
| MYC23(L4W) | EQK**W**ISEEDL |
| MYC24(L4Y) | EQK**Y**ISEEDL |
| MYC27(I5A) | EQKL**A**SEEDL |

| | |
|---|---|
| MYC28(I5C) | EQKL**C**SEEDL |
| MYC29(I5D) | EQKL**D**SEEDL |
| MYC30(I5E) | EQKL**E**SEEDL |
| MYC31(I5F) | EQKL**F**SEEDL |
| MYC32(I5H) | EQKL**H**SEEDL |
| MYC33(I5K) | EQKL**K**SEEDL |
| MYC34(I5L) | EQKL**L**SEEDL |
| MYC35(I5M) | EQKL**M**SEEDL |
| MYC36(I5N) | EQKL**N**SEEDL |
| MYC37(I5P) | EQKL**P**SEEDL |
| MYC38(I5Q) | EQKL**Q**SEEDL |
| MYC39(I5R) | EQKL**R**SEEDL |
| MYC40(I5S) | EQKL**S**SEEDL |
| MYC41(I5T) | EQKL**T**SEEDL |
| MYC42(I5V) | EQKL**V**SEEDL |

**# the replaced residues were highlighted in red color.**

**Supplementary Table S4 Sequences of Flag-tag and its mutants**

| **Number** | **Sequences** |
|---|---|
| **Flag**-tag | **DYKDDDDK** |
| Flag01(K3G) | DY**G**DDDDK |
| Flag02(K8G) | DYKDDDD**G** |
| Flag03(K3A) | DY**A**DDDDK |
| Flag04(K3C) | DY**C**DDDDK |
| Flag05(K3D) | DY**D**DDDDK |
| Flag06(K3E) | DY**E**DDDDK |
| Flag07(K3F) | DY**F**DDDDK |
| Flag08(K3H) | DY**H**DDDDK |
| Flag09(K3I) | DY**I**DDDDK |

| | |
|---|---|
| Flag10(K3L) | DY**L**DDDDK |
| Flag11(K3M) | DY**M**DDDDK |
| Flag12(K3N) | DY**N**DDDDK |
| Flag13(K3P) | DY**P**DDDDK |
| Flag14(K3Q) | DY**Q**DDDDK |
| Flag15(K3R) | DY**R**DDDDK |
| Flag16(K3S) | DY**S**DDDDK |
| Flag17(K3T) | DY**T**DDDDK |
| Flag18(K3V) | DY**V**DDDDK |
| Flag19(K3W) | DY**W**DDDDK |
| Flag20(K3Y) | DY**Y**DDDDK |

**# the replaced residues were highlighted in red color.**

**Supplementary Table S5 Sequences of VSV-tag and its mutants**

| **Number** | **Sequences** |
|---|---|
| VSV-tag | **YTDIEMNRLGK** |
| VSV01(T2G) | Y**G**DIEMNRLGK |
| VSV02(I4G) | YTD**G**EMNRLGK |
| VSV03(M6G) | YTDIE**G**NRLGK |
| VSV04(N7G) | YTDIEM**G**RLGK |
| VSV05(R8G) | YTDIEMN**G**LGK |
| VSV06(T2A) | Y**A**DIEMNRLGK |
| VSV07(T2C) | Y**C**DIEMNRLGK |
| VSV08(T2D) | Y**D**DIEMNRLGK |
| VSV09(T2E) | Y**E**DIEMNRLGK |
| VSV10(T2F) | Y**F**DIEMNRLGK |
| VSV11(T2H) | Y**H**DIEMNRLGK |
| VSV12(T2I) | Y**I**DIEMNRLGK |
| VSV13(T2K) | Y**K**DIEMNRLGK |

| | |
|---|---|
| VSV14(T2L) | Y**L**DIEMNRLGK |
| VSV15(T2M) | Y**M**DIEMNRLGK |
| VSV16(T2N) | Y**N**DIEMNRLGK |
| VSV17(T2P) | Y**P**DIEMNRLGK |
| VSV18(T2Q) | Y**Q**DIEMNRLGK |
| VSV19(T2R) | Y**R**DIEMNRLGK |
| VSV20(T2S) | Y**S**DIEMNRLGK |
| VSV21(T2V) | Y**V**DIEMNRLGK |
| VSV22(T2W) | Y**W**DIEMNRLGK |
| VSV23(T2Y) | Y**Y**DIEMNRLGK |
| VSV26(M6A) | YTDIE**A**NRLGK |
| VSV27(M6C) | YTDIE**C**NRLGK |
| VSV28(M6D) | YTDIE**D**NRLGK |
| VSV29(M6E) | YTDIE**E**NRLGK |
| VSV30(M6F) | YTDIE**F**NRLGK |
| VSV31(M6H) | YTDIE**H**NRLGK |
| VSV32(M6I) | YTDIE**I**NRLGK |
| VSV33(M6K) | YTDIE**K**NRLGK |
| VSV34(M6L) | YTDIE**L**NRLGK |
| VSV35(M6N) | YTDIE**N**NRLGK |
| VSV36(M6P) | YTDIE**P**NRLGK |
| VSV37(M6Q) | YTDIE**Q**NRLGK |
| VSV38(M6R) | YTDIE**R**NRLGK |
| VSV39(M6S) | YTDIE**S**NRLGK |
| VSV40(M6T) | YTDIE**T**NRLGK |
| VSV41(M6V) | YTDIE**V**NRLGK |
| VSV42(M6W) | YTDIE**W**NRLGK |
| VSV43(M6Y) | YTDIE**Y**NRLGK |
| VSV35(M6N) | YTDIE**N**NRLGK |

| | |
|---|---|
| VSV36(M6P) | YTDIE**P**NRLGK |
| VSV37(M6Q) | YTDIE**Q**NRLGK |
| VSV38(M6R) | YTDIE**R**NRLGK |
| VSV39(M6S) | YTDIE**S**NRLGK |
| VSV40(M6T) | YTDIE**T**NRLGK |
| VSV41(M6V) | YTDIE**V**NRLGK |
| VSV42(M6W) | YTDIE**W**NRLGK |
| VSV43(M6Y) | YTDIE**Y**NRLGK |

**# the replaced residues were highlighted in red color.**
